# Polar and non-polar fractions of deep fried edible oils induce differential cytotoxicity and hemolysis

**DOI:** 10.1101/2021.02.03.429519

**Authors:** P. Sneha, Yemeema Paul, Mithula Venugopal, Arunaksharan Narayanankutty

## Abstract

Edible oils are the essential part of diet, however, deep frying process induce oxidative changes in these oils, making them unsuitable for consumption. Deep frying generates various noxious polar and non-polar aldehydes and carbonyls, which may be polar or non-polar in nature. The present study thus evaluated the cytotoxic and hemolytic effects of polar and non-polar fractions of different deep fried edible oils. There observed a significantly elevated level of lipid peroxidation products in the polar fraction of deep fried sunflower (FSO-P) and rice bran oils (FRO-P). The treatment with these fractions induced cytotoxicity in cultured colon epithelial cells, with a higher intensity in FSO-P and FRO-P. Further, an increased TBARS level and catalase activity in RBCs treated with FSO-P and FRO-P led to hemolysis. In comparison, the fried coconut oil (FCO) fractions were less toxic and hemolytic; in addition, the non-polar fraction was more toxic, compared to FCO-P fraction.

## 1. Introduction

Edible oils form the essential part of daily diet, which are providing the essential fatty acids, certain vitamins, and other bioactive compounds. Chemically these oils are triglycerides, where the nature of fatty acids attached (saturated or unsaturated) determine the chemical, physical and health properties of the oil. The cuisine systems use various methods for the use of edible oils, of which deep frying is the prominent.

The deep frying exposes these edible oils to high temperature and oxygen, which accelerates the oxidative modifications in the oil. A study by Choe and Min (2007) and Warner (1999) have classified the deep frying induced changes in to oxidation, hydrolysis and polymerization. Predominantly, the hydrolysis and oxidation are taking place in the edible oils with high unsaturation, making them unhealthy for consumption. However, the triglyceride polymerization can take place in saturated as well as unsaturated edible oils. The noxious products formed during the deep frying products are reported to be unhealthy due to several reasons. A volume of studies have identified a positive association between fried oil intake and hypertension (Kamisah et al. 2016, Kamisah et al. 2015, Leong et al. 2010). In a cross sectional anthropometric study, there observed increased association between hypertension and intake of thermally oxidized sunflower oil, especially that is rich in polar compounds (Soriguer et al. 2003). Corroborating with these, consumption of repeatedly heated soybean and palm oil increases the VCAM-1 and ICAM levels in rats (Ng et al. 2012a, Ng et al. 2012b). Hence, it can be ascertained that the alterations in the vascular thickening and vascular inflammation leads to hypertensive disorders during thermally oxidized edible oil feeding. Consumption of repeatedly heated coconut oil induce hepatic foci and pre-neoplastic lesions in rats treated with diethyl nitrosamine (Srivastava et al. 2010a). Similarly to these, boiled sunflower and mustard oil are shown to have genotoxic and carcinogenic effects in murine models (Shukla and Arora 2003, Srivastava et al. 2010b).

Some of the compounds identified include Trans-trans-2,4-decadienal, a derivative during frying of peanut oil, is shown to induce genotoxicity mediated by the formation of reactive oxygen species and reduction of cellular glutathione content (Wu and Yen 2004). 1-nitropyrene and 1,3-dinitropyrene are the derivatives of lard, soybean and peanut oils (Wu et al. 1998).

The chemical nature of the toxic compounds formed is still not clear; it has been identified that the non-polar fractions of deep fried coconut oil induce hepatotoxicity and lipotoxicity in animals (Narayanankutty et al. 2018). However, some studies have indicated that polar fractions isolated are inducing deleterious effects such as in peanut oil (Ju et al. 2019, Li et al. 2016). However, most of these studies failed to compare the toxic effects of both polar and non-polar fractions. The present thus aims to provide a clear role of the polar and non-polar fractions of different edible oils on cytotoxicity and hemolysis. In addition, emphasize is given on the redox status of the cells during their cytotoxic effects.

## 2. Materials and Methods

### 2.1 Edible oils used in the study

Coconut oil, sunflower oil, and rice bran oil used for deep frying of chips were collected from commercial chips manufacturers, whose identity is not disclosed. The oils were stored in amber colored bottles under -20°C.

### 2.2 Isolation of polar and non-polar fractions

Total polar and non-polar contents of these oils were isolated by the column chromatography. The glass column was packed with 25 g of silica gel (200-400 mesh) and washed three times with petroleum ether/diethyl ether (87:13, v/v). The column was loaded with 5 g of each fried oils together with petroleum ether/diethyl ether (87:13, v/v). The nonpolar compounds were eluted with 150 mL of the aforementioned solvent system; the polar fraction was eluted with 150 mL of diethyl ether. The eluent was dried and stored under -20°C. The concentrate was further dissolved in tetrahydrofuran (THF) for further analysis.

### 2.3 Biochemical analysis

Changes in the lipid peroxidation indicators such as thiobarbituric acid reactive substances (TBARS) (Pegg 2004), conjugated diene (CD) as well as conjugated triene (CT) (Pegg 2004) were estimated as per the standard protocol.

### 2.4 Cytotoxicity and hemolysis assay

The human colon epithelial cells (HCT-116) were cultured in a 24 well plate at a density of 1×10^6^ cells/ mL. After 24 hours, the isolated polar and non-polar fractions of different edible oils were added to each well. A set of wells were maintained as control and THF control; it was further incubated for 48 hours. At the end of incubation, MTT was added to each well mixed and allowed to develop formazan crystals for 4 hours (Mosmann 1983). The crystals were dissolved in DMSO and absorbance was read at 570 nm. The percentage cell death was calculated using the formula;

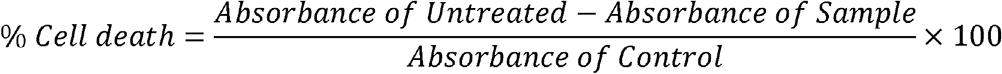

### 2.5 Hemolysis assay

The blood was collected from slaughter house in a tube coated with anti-coagulant; the blood was washed with physiological saline for three times and RBCs were pelleted out. The blood pellet was further diluted to a 23% with Phosphate buffered saline.

To test the hemolytic effects, of blood (23%) was mixed with 100 μL of polar and non-polar fractions of different edible oils (0-250 μg/mL). The mixture was then incubated at 37° C for 1 hour and then diluted with PBS (2.4 mL). The blood sample was centrifuged at 3000g for 20 mins and the absorbance of the supernatant was measured at 450 nm. The percentage increase in hemolysis was estimated by comparing with the control. The pellet of centrifugation was used to quantify the lipid peroxidation products as thiobarbituric acid reactive substances (Ohkawa et al. 1979) as well as catalase activity (Beers and Sizer 1952).

### 2.6 Statistical analysis

The values are expressed as mean± SD of three independent experiments, each carried out in triplicate. The statistical analysis was done using one way ANOVA followed by Tukey Kramer multiple comparison test.

## 3. Results and Discussion

Lipid peroxidation products are recognized as a driving force in hypertension (Jaarin et al. 2011), dyslipidemia and oxidative stress (Adam et al. 2008, Liu et al. 2014), impaired glycerolipid metabolism as well as gut microbiota (Zhou et al. 2016),.non-alcoholic fatty liver (Konishi et al. 2006, Narayanankutty et al. 2017), and cancers (Skrzydlewska et al. 2005). Together with this association of fried foods can be observed with lifestyle associated diseases (Gadiraju et al. 2015).

The lipid peroxidation products including Conjugated diene (CD), conjugated triene (CT) and thiobarbituric acid reactive substance (TBARS) are found to be elevated in the non-polar fraction of fried coconut and rice bran oils (Table 1). On contrary, in deep fried sunflower oil, the polar fraction had higher lipid peroxidation products in comparison with the non-polar fraction. Among the different edible oils, the quantity of lipid peroxidation products were high in the FSO polar fraction was predominant. It has already been proven that the polyunsaturated fat rich edible oils undergo rapid oxidative modifications during deep frying. Therefore, the high linoleic acid content of FSO might be the reason for elevated level of lipid peroxidation products in FSO. Apart from that, it has been reported that the deep frying and thermal oxidation results in the increased polar compounds in edible oils; therefore, the polar lipid peroxides and carbonyls formed during the deep frying of sunflower oil is expected to the observed increase in lipid peroxidation products. This is in accordance with the previous studies conducted by Moros et al. (2009) where similar increase in lipid peroxidation products are reported. Together with the oxidation products, a study have reported the formation of cyclic fatty acid monomers (CFAM), in unsaturated edible oils (Romero et al. 2006). These CFAMs are known to be deleterious to the body and has been proven to have enzyme inhibitory properties (Lamboni et al. 1998).

**Table 1.** Changes in the biochemical parameters in polar and non-polar fractions of fried oils

| Edible oil | Conjugated<br>dienes | Conjugated<br>trienes | TBARS |
| --- | --- | --- | --- |
| FCO-P | 0.224±0.011 | 0.102±0.008 | 0.443±0.062 |
| FCO-N | 0.323±0.017** | 0.203±0.017** | 0.688±0.085** |
| FRO-P | 0.299±0.020 | 0.467±0.020 | 0.945±0.095 |
| FRO-N | 0.313±0.022* | 0.527±0.024** | 1.062±0.106** |
| FSO-P | 0.488±0.021 | 0.716±0.028 | 1.528±0.173 |
| FSO-N | 0.359±0.024** | 0.556±0.039** | 0.983±0.118*** |

FCO- Fried coconut oil, FRO- Fried rice bran oil, FSO- Fried sunflower oil, and the abbreviations P- polar and N- non-polar fractions; The values are represented as mean± SD of three independent experiments, each carried in triplicate. (\* indicate p<0.05; \*\* indicate p<0.01; \*\*\* indicate p<0.001)

The cytotoxicity of the polar and non-polar fractions of different oils were evaluated in human colon cancer cells. There observed a significantly higher toxicity (IC50 value 311.72± 19.2 µg/mL) in polar fraction of FSO (p<0.01) compared to the polar and non-polar fractions of other oils. Similarly, the FRO-P fraction had a marginally higher cytotoxicity (475.29±11.4 µg/mL) than its non-polar fraction (488.28±16.2 µg/mL). On contrary, the cytotoxic effect of non-polar fraction of FCO was higher (592.42± 13.7 µg/mL) compared to its polar fraction (765.7± 24.9 µg/mL). Previous study by Ju et al. (2019) observed a similar cytotoxic effect of polar fractions derived from the deep fried peanut oil, where the cytotoxicity was mediated by disruption of antioxidant system and also by inducing cell cycle arrest. Further studies reported that the oxidized triglycerides are responsible for the cytotoxic effect of the polar fraction (Li et al. 2016). In addition, the probable role of an oleic acid oxidation product, Epoxy stearic acid, on the cytotoxic effect is also identified (Liu et al. 2018). It is thus possible that the toxic effects of different polar fractions of deep fried oils observed in the present study may also be mediated through redox imbalance.

In connection with the cytotoxic properties, the hemolytic properties of the different fractions in RBC were also evaluated. The results were also similar to that of cytotoxic effect, with the highest hemolytic potential in the FSO-P fraction (EC50 value 59.4± 4.5 µg/mL). The polar fraction of FRO (EC50= 69.0± 7.9 µg/mL) and non-polar fraction of FCO (EC50= 117.1± 12.2 µg/mL) were more hemolytic in the respective oil groups. Among these, the polar fraction of FCO was least hemolytic in nature (EC50= 153.0± 17.5µg/mL). It has been reported that linoleic acid and arachidonic acid content can exacerbate the oxidative hemolysis of erythrocytes (Yuan et al. 2015, Yuan et al. 2017). It is therefore possible that the linoleic acid contents and their oxidation products in FSO-P may be responsible for the increased oxidative hemolysis. Apart from this, it has been reported that the intrinsic lipid peroxidation status and antioxidant enzyme activities determine the hemolytic process (Sudha et al. 2004, Singh and Rahman 1987). Supporting their observations, the results of the study also observed significant alterations in the redox status of erythrocytes.

As indicated in Table 2, the level of thiobarbituric acid reactive substances were significantly higher in the erythrocytes treated with the polar fraction of FSO (p<0.01) and FRO (p<0.05). The lipid peroxidation status in FCO-P treated erythrocytes was marginally higher, but without statistical significance. Together with the TBARS levels, the catalase activity in the FSO-P treated erythrocytes were significantly high (p<0.01); similarly, a significant increase in the catalase activity was also observed with all other treatment groups, with the exception in FCO-P. It is expected that the peroxides and aldehydes in the deep fried oils, especially FSO-P fraction, may have increased the membrane lipid peroxidation and subsequently lead to hemolysis (Sudha et al. 2004). Corroborating with this, there observed significant elevation in the catalase activity in FSO-P and FRO-P fractions. It has been reported that catalase activity is positively correlated to the cellular peroxide insults (Meilhac et al. 2000).

**Table 2.** Levels of various antioxidant parameters in the erythrocytes treated with different fractions of edible oils

| **Edible oil** | **Catalase**<br>**(U/mg protein)** | **TBARS**<br>**(nmoles/mg protein)** |
| --- | --- | --- |
| Untreated | 56.82±4.7 | 3.66±0.44 |
| FCO-P | 67.93±7.2<sup>ns</sup> | 4.12±0.28<sup>ns</sup> |
| FCO-N | 74.50±6.1* | 4.83±0.66* |
| FRO-P | 86.11±9.5* | 5.60±0.49** |
| FRO-N | 77.84±5.8* | 4.47±0.78* |
| FSO-P | 93.45±7.6** | 6.77±0.71** |
| FSO-N | 81.29±6.4* | 5.17±0.65* |

FCO- Fried coconut oil, FRO- Fried rice bran oil, FSO- Fried sunflower oil, and the abbreviations P- polar and N- non-polar fractions; The values are represented as mean± SD of three independent experiments, each carried in triplicate. (\*indicate significant difference p<0.05; \*\* indicate significant difference p<0.01)

It is therefore possible that the toxic effects of deep fried edible oils of polyunsaturated fat containing edible oils are mediate through its polar fraction. However, in saturated fat containing edible oils, the toxic compounds are present in the non-polar fraction. Previously, a study has indicated that hepatotoxic effect and lipotoxicity of deep fried coconut oil is mediated through its non-polar fraction (Narayanankutty et al. 2018).

## 4. Conclusion

The study concludes that deep fried oil polar and non-polar fractions induce oxidative hemolysis and cytotoxicity. In polyunsaturated fatty acid containing edible oils, the toxic components are predominantly present in the polar fraction; however, in saturated fat rich deep fried oils, the non-polar fraction is more toxic. It is possible that the differential nature of the peroxides formed during deep frying of polyunsaturated and saturated fat containing edible oils contributed to the differentials effects. However, the study doesn’t rule out the possible toxic effects of other fractions, only the results indicate a potentially higher toxicity, thereby indicating the possible extraction of toxic substance to that fraction. Therefore, the study recommends reducing the use of deep fried food items.

## Acknowledgement

Authors acknowledge the financial support from Kerala State Council for Science, Technology, and Environment in the form of student project scheme.

---

**Figure 1.**
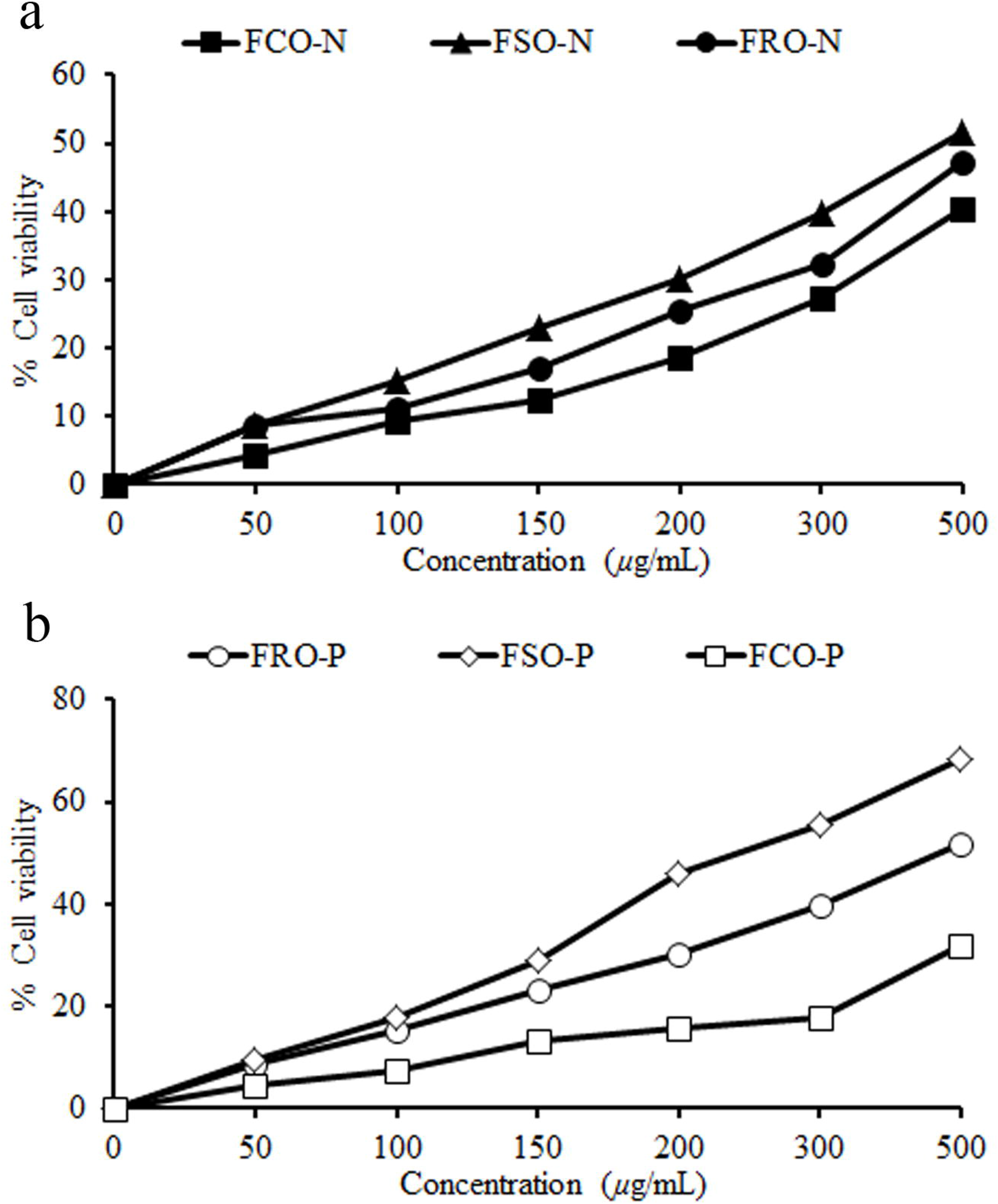
The cytotoxic effect of polar (P) and non-polar (N) fractions of deep fried coconut (FCO), rice bran (FRO) and sunflower oil (FSO) in human colon epithelial cells.

**Figure 2.**
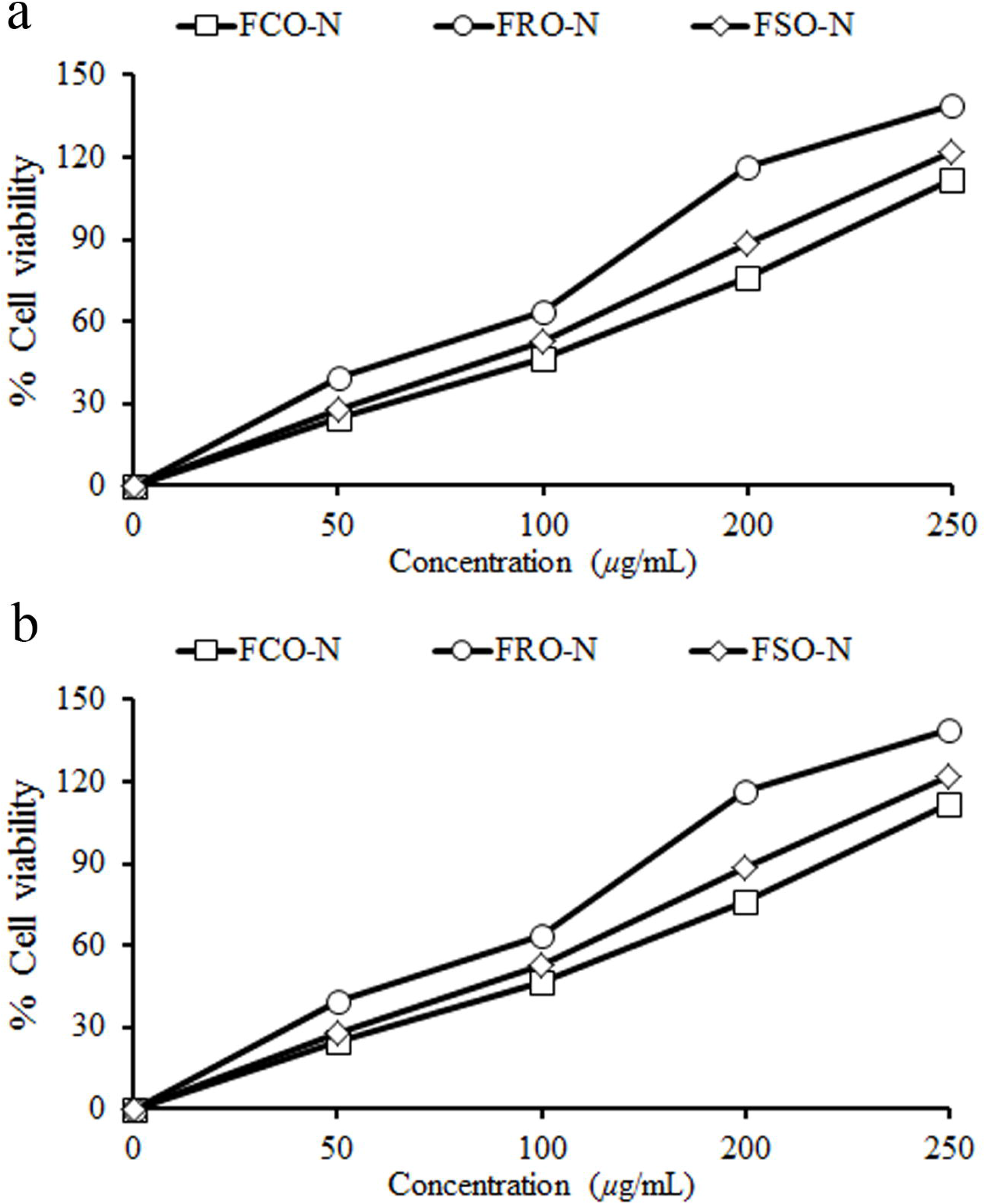
The hemolytic properties of the polar (P) and non-polar (N) fractions of deep fried coconut (FCO), rice bran (FRO) and sunflower oil (FSO) in erythrocytes.

